# An amplicon-based sequencing framework for accurately measuring intrahost virus diversity using PrimalSeq and iVar

**DOI:** 10.1101/383513

**Authors:** Nathan D Grubaugh, Karthik Gangavarapu, Joshua Quick, Nathaniel L. Matteson, Jaqueline Goes De Jesus, Bradley J Main, Amanda L Tan, Lauren M Paul, Doug E Brackney, Saran Grewal, Nikos Gurfield, Koen KA Van Rompay, Sharon Isern, Scott F Michael, Lark L Coffey, Nicholas J Loman, Kristian G Andersen

## Abstract

How viruses evolve within hosts can dictate infection outcomes; however, reconstructing this process is challenging. We evaluated our multiplexed amplicon approach - PrimalSeq - to demonstrate how virus concentration, sequencing coverage, primer mismatches, and replicates influence the accuracy of measuring intrahost virus diversity. We developed an experimental protocol and computational tool (iVar) for using PrimalSeq to measure virus diversity using Illumina and compared the results to Oxford Nanopore sequencing. We demonstrate the utility of PrimalSeq by measuring Zika and West Nile virus diversity from varied sample types and show that the accumulation of genetic diversity is influenced by experimental and biological systems.

## Background

RNA viruses, including HIV, influenza, West Nile, and Zika, pose significant threats to public health worldwide. Part of this burden stems from their ability to rapidly evolve within hosts [1]. Generation of intrahost genetic diversity allows virus populations to evade host immune responses [2–4], alter the severity of disease [5], and adapt to changing environments [6,7]. Studying virus populations, both within naturally infected hosts and during experimental evolution, can therefore lead to breakthroughs in our understanding of virus-host interactions and novel approaches for outbreak response [8–11].

In many cases, however, accurately measuring intrahost RNA virus diversity using deep sequencing remains a significant challenge. Multiple factors, such as virus titer, sample preparation, sequencing errors, and computational inferences, can bias measures of genetic diversity [12–16]. Moreover, for many clinical samples, low ratios of viral to host RNA often necessitates enrichment of viral nucleic acid to recover sufficient templates for deep sequencing [17]. This is especially true for Zika virus, where low viremias (<1000 copies/μL of RNA) are often detected during natural and experimental infections [18–21]. PCR amplification of virus nucleic acid is a common approach to overcome this challenge [4,22,23], although it can introduce biases by altering the composition of intrahost genetic variants [14,24]. Therefore, to ensure accuracy, comprehensive validation of deep sequencing approaches should accompany diversity measures from biological samples.

We previously developed a multiplex primer design tool (‘Primal Scheme’) coupled to a laboratory protocol (‘PrimalSeq’) to sequence RNA viruses directly from clinical samples in a way that is cheap, accurate, and scalable under resource-limited conditions [17]. Versions of PrimalSeq have been used to sequence the majority of Zika virus genomes from the epidemic in the Americas [19,25–27], yellow fever virus in Brazil [28], and West Nile virus in the United States [29]. PrimalSeq has also been used to characterize Zika virus during infection of non-human primates [21,30]. While PrimalSeq was shown to be superior to other methods for obtaining consensus sequences [25,26], it has yet to be validated for measuring intrahost diversity.

In this study, we benchmarked PrimalSeq for sequencing diverse virus populations, highlighting its limitations and providing recommendations for accurately measuring intrahost single nucleotide variants (iSNVs) from both Illumina and Oxford Nanopore data. We used these results to develop comprehensive laboratory protocols and a computational tool (iVar), and further tested PrimalSeq to characterize Zika virus populations generated from cell culture, mosquito, non-human primate, and human clinical samples. We demonstrate the utility of PrimalSeq for other viruses by designing an amplicon scheme for West Nile virus and measuring genetic diversity from field collected mosquitoes and birds. Our data show that virus diversity can be significantly impacted by the experimental and biological systems, and we provide a framework to uncover the underlying mechanisms. PrimalSeq and iVar provide a scalable platform for viruses other than Zika and West Nile that can be applied to discover ecological, epidemiological, and immunological drivers of virus evolution in a variety of systems.

## Results

### Virus concentration and sequencing depth impact intrahost variant calling

We previously developed Primal Scheme (Quick *et al.* - primal.zibraproject.org), a multiplex primer design tool for amplicon-based sequencing of RNA virus genomes directly from clinical samples [17]. Our Zika virus PrimalSeq protocol generates 35 overlapping amplicons of ~400 base pairs from two multiplexed PCR reactions, an approach similar to ‘RNA jackhammering’, which was developed to sequence HIV [31]. The process of PCR amplification to generate sufficient templates for high-throughput sequencing, however, may bias the measurements of intrahost virus diversity through differential amplification efficiencies for divergent virus haplotypes present in a population [32,33].

Given the potential biases that may be introduced during PCR amplification, we sought to assess the accuracy of PrimalSeq for iSNV detection. Through a series of experiments, we determined that at least 1000 RNA virus copies are needed to detect iSNVs at greater than 3% frequency, when sequenced to a coverage depth of at least 400× (*i.e.*, the number of sequenced nucleotides per targeted genome position, **Fig. 1**). To set up these experiments, we created genetically diverse Zika virus populations by mixing two divergent virus strains: Zika virus #1 isolated from Puerto Rico in 2015 (Genbank KX087101) and virus #2 isolated from Cambodia in 2010 (Genbank KU955593). Using gold-standard untargeted metagenomic sequencing [34], we determined that there were 159 fixed consensus sequence differences between the viruses located throughout the 10,807 nucleotide genome of virus #1 and #2 (**Fig. 1a**).

**Figure 1.**
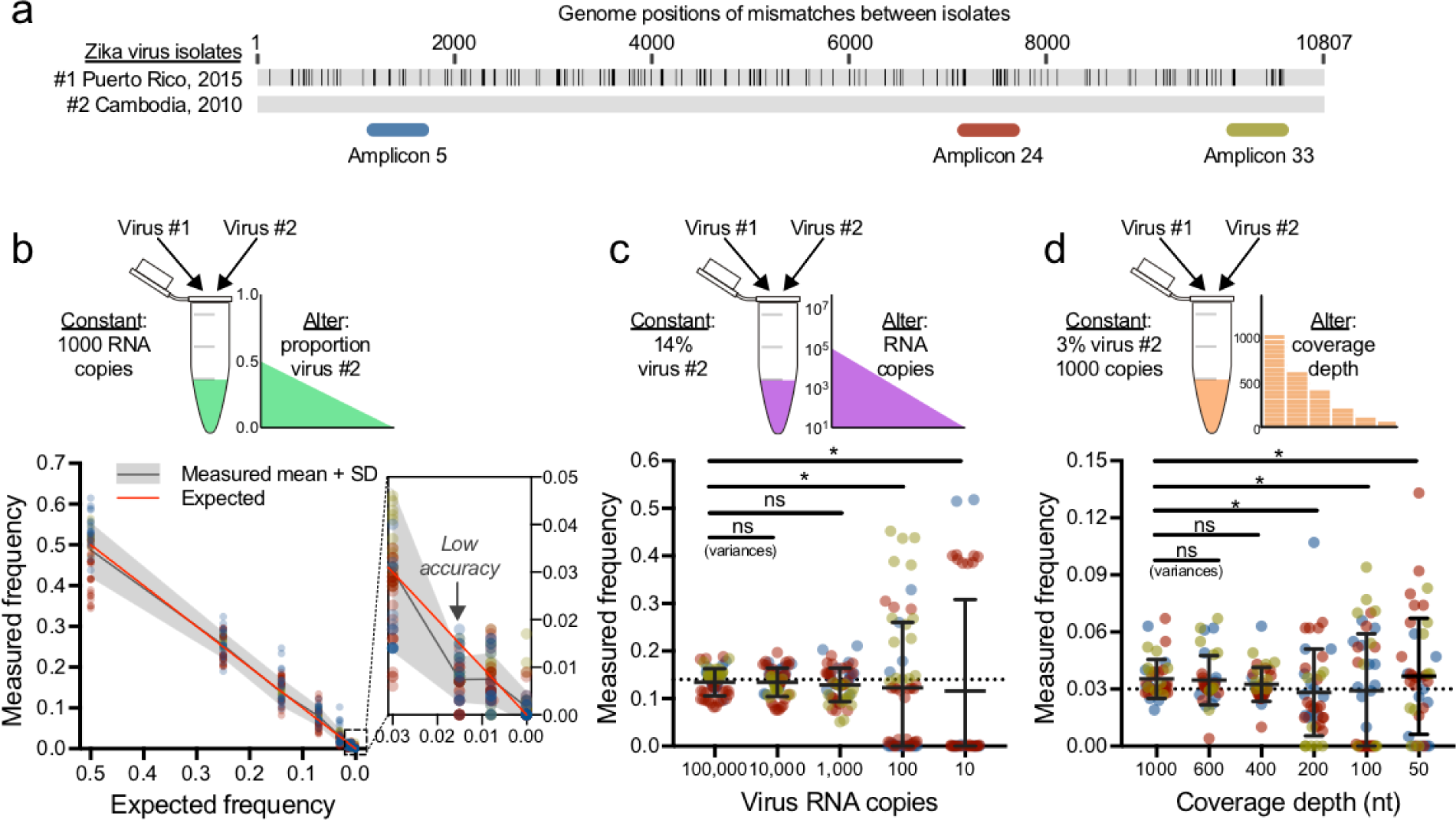
Measurement intrahost variant frequencies are more accurate at high frequencies and are susceptible to input concentrations and coverage depths. (**a**) We created genetically diverse virus populations by mixing two Zika virus isolates with 159 consensus nucleotide differences to test the effects of PCR amplification prior to sequencing to measure intrahost single nucleotide variant (iSNV) frequencies. For these initial experiments, we amplified three ~400 bp regions of the Zika virus genome using primers without any mismatches to either of the mixed virus (shown as amplicons 5, 24, and 33). Amplicon 5 contains five iSNV sites, amplicon 24 contains 8 iSNV sites, and amplicon 33 contains 5 iSNV sites. (**b**) We created virus populations containing 50%, 25%, 14%, 7%, 3%, 1.5%, and 0.8% virus #2 to test the impact of PCR amplification prior to sequencing on measuring ranges of iSNV frequencies. The data points represent individual iSNVs amplified and sequenced in triplicate from each population (colored by amplicon 5, 24, or 33 as shown in Fig. 1a). (**c**) We 10-fold serially diluted a mixed population containing 14% of virus #2 (expected, dotted line) from 100,000 to 10 copies to test the effects of input concentrations on accurate iSNV measurements. (**d**) We randomly downsampled the datasets generated from 1000 input virus RNA copies containing 3% virus #2 to set coverage depths (sequenced nucleotides [nt] per genome position) to determine the minimum coverage needed to yield accurate iSNV measurements. For **c** and **d**, the Levene’s test was used to assess equality among variances of iSNV measurements from each coverage depth (ns, not significant; *, *p* < 0.05). Data shown as means with standard deviations.

For our initial evaluation, we selected three of the 35 Zika virus primer sets that flanked at least five variable genome positions (amplicons 5, 24, and 33). We then made two sets of mixed virus populations: (***1***) altering the ratios of mixed viruses, while keeping the overall input concentration constant at 1000 virus RNA copies (**Fig. 1b**), and (***2***) maintaining a constant ratio of 14% of virus #2 while altering the input concentrations of virus RNA used for cDNA synthesis (**Fig. 1c**). For each test, we measured the frequencies of the 19 iSNVs between virus #1 and #2 (**Fig. 1a**). We generated the amplicons independently three times and sequenced each using the Illumina MiSeq platform (‘technical replicates’).

We found that the measured mean iSNV frequencies were accurate from populations containing 50%, 25%, 14%, 7%, and 3% of virus #2 (**Fig. 1b**). At 1.5% of virus #2, the standard deviation of our measured mean iSNV frequency (0.2%-1.2%) fell below the expected frequency (**Fig. 1b**). This demonstrates that the lower limit of accurate iSNV detection for PrimalSeq in this scenario is between 1.5-3%. When we altered input concentrations of a population containing 14% virus #2 from 100,000 to 10 virus RNA copies (10-fold serial dilutions), we found that the variances of measured frequencies became significantly higher from concentrations containing 100 or less copies (Levene’s test for variance, *p* < 0.05; **Fig. 1c**). Therefore, input virus concentrations can dramatically alter iSNV detection, as others have also discovered [12]. We conclude that a minimum of 1000 virus RNA copies should be used with PrimalSeq to accurately measure iSNVs greater than 3% frequency.

Sequencing coverage depth is another important factor for iSNV detection [13], so we sought to define the level of sequencing coverage needed to accurately measure iSNVs. From our samples containing 1000 virus RNA copies with 97% virus #1 and 3% virus #2, we sequenced the targeted genome regions in triplicates to a coverage depth of ~3000×. We randomly downsampled these datasets to generate coverage depths of 1000, 600, 400, 200, 100, and 50× (**Fig. 1d**). We found that the variances of iSNV frequencies became significantly higher from coverage depths lower than 400× (*i.e.*, at 200, 100, and 50×; Levene’s test for variance, *p* < 0.05; **Fig. 1d**). Thus, we conclude that a minimum sequencing coverage depth of 400× is required to maintain iSNV measurement accuracy at the lower limit of frequency detection (3%) and input concentration (1000 virus RNA copies).

### Primer mismatches impact intrahost variant frequency measurements

A concern for using PCR-based sequencing protocols for measuring intrahost virus diversity is the potential for primer mismatches to alter PCR efficiency that can bias iSNV frequency measurements. Indeed, we found that primer mismatches, especially those close to the 3’ end, can alter iSNV frequencies in the generated amplicons. To make this assessment, we used (***1***) our Zika virus PrimalSeq strategy and (***2***) a mix containing 90% of virus #1 and 10% of virus #2 (‘Mix10%’). Of the 159 consensus nucleotide differences between virus #1 and #2, 24 resulted in mismatches within the primer regions of 20 of the oligos used to generate 18 of the 35 PCR amplicons (**Fig. 2a**, **Table S1**). In addition, at least one nucleotide difference occured within each of the 18 amplicons (outside of the primer-binding region), which allowed us to assess the influence of primer mismatches on iSNV frequency measurements across the Zika virus genome.

**Figure 2.**
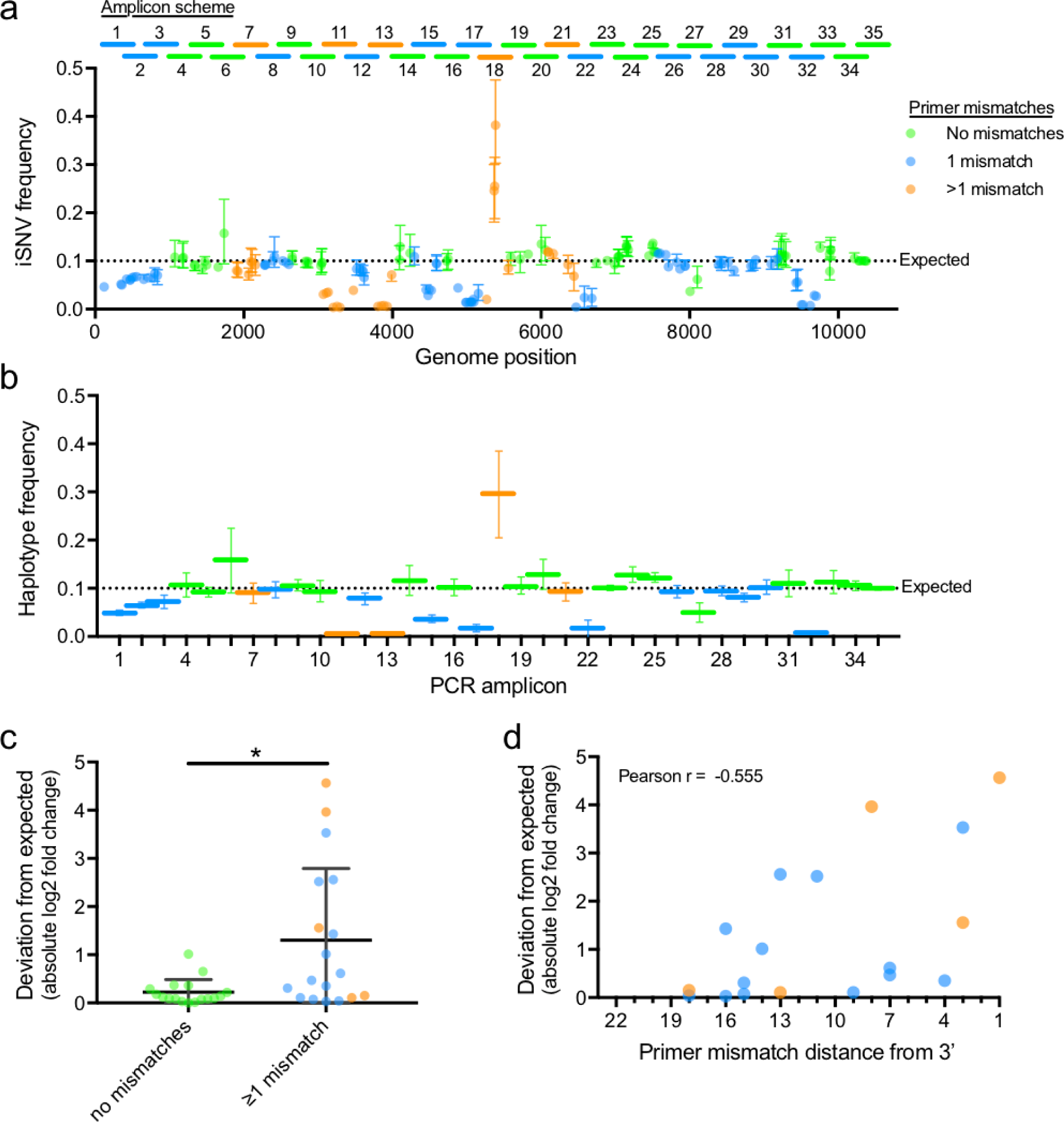
Measures of intrahost variant frequencies are sensitive to primer mismatches. (**a**) To assess the impacts of primer mismatches on accurately measuring intrahost single nucleotide variants (iSNVs), we sequenced a mixed Zika virus population using 35 overlapping PCR amplicons (see “Amplicon scheme” above panel). The virus population contained 10% virus #2 (Expected) and 1000 virus RNA copies were amplified and sequenced in triplicate. The amplicons and iSNVs are colored according to the number of mismatches in the primer sequences used to generate that amplicon. Data shown as means and ranges. (**b**) To account for unequal iSNV sites within each amplicon, the iSNV frequencies on each amplicon were averaged to produce a haplotype frequency for virus #2 mixed at 10% (Expected). Data shown as means and ranges. (**c**) We calculated the deviations between the measured and expected virus #2 haplotype frequencies (absolute value of the log2 fold change) to assess the bias introduced during PCR of amplicons containing primer mismatches to virus #2 (*, Welch’s t test, *p* < 0.05). Data shown as means and standard deviations. (**d**) We plotted the deviations from expected haplotype frequencies by the distance of mismatches from the 3’ end of the primer to investigate the impact of mismatch location. If more that one mismatch was present on a primer pair (orange), the data is shown using the closest mismatch to the 3’ end. Mismatches closer to the 3’ end of the primer are more likely decrease the accuracy of iSNV or haplotype measurements from that amplicon (correlation by Pearson r, *p* < 0.05). Data shown as the mean from all three replicates.

We amplified the Mix10% virus population (1000 RNA copies) independently three times and sequenced each replicate to a minimum coverage depth of 1,000× using the Illumina MiSeq. We measured iSNV frequency at each site and calculated the mean iSNV frequency of all iSNVs within an amplicon to estimate the computed virus #2 haplotype frequency (**Fig. 2b**). We found that iSNV frequencies measured from amplicons without primer mismatches were significantly closer to the expected value of 10%, than amplicons with one or more mismatches in the primer regions (Welch’s t test, *p* < 0.05, **Fig. 2c**). Moreover, we found that mismatches closer to the 3’ end of the primer were more likely to lead to inaccurate frequency measurements (Pearson r, *p* < 0.05, **Fig. 2d**). Overall, our data demonstrate that the accuracy of intrahost virus diversity measures is highly impacted by primer mismatches during PCR. Thus, when iSNVs are detected from amplicons with mismatches in the primer binding sites, the resulting diversity data from those amplicons should be interpreted with caution.

### Removal of false positives intrahost variants with replicate sequencing

Measurements of virus intrahost genetic diversity are sensitive to PCR and sequencing errors [12,14,16]. These factors, combined with others such as virus concentration and sequencing coverage (**Fig. 1**), can lead to erroneous iSNV detection (*i.e.*, false positives) and bias measures of genetic diversity. To improve accurate iSNV detection, we examined the distribution of false positive iSNV calls and investigated methods to remove them during analysis. We found that (***1***) the distribution of false positive iSNVs more closely matched the profile of sequencing errors than PCR errors, and that (***2***) the majority of false iSNV >3% could be removed by replicate sequencing.

To investigate false positive iSNV calls, we amplified our Mix10% virus population (1000 RNA copies) individually three times and sequenced each on the Illumina MiSeq. We limited our analysis to only those amplicons with perfect primer matches. Within these regions, we analyzed 54 sites with expected 10% iSNVs (true positives) and 4173 sites that were invariable in our mixed virus population. We considered any iSNVs detected >0.1% frequency at the invariable sites as false positives. We found that on average, 631 of the 4173 expected invariable sites (16%) had false positive iSNVs at a >0.1% frequency cut-off. We observed false positive iSNVs on every amplicon but found that they were unevenly distributed across the 250 nucleotide long Illumina reads (**Fig. 3a-b**). Specifically, we found that virus genome sites covered by sequencing reads at positions >150 nucleotides had significantly more false positives (Wilcoxon test, *p*< 0.05; **Fig. 3a** inset) at significantly higher frequencies (Mann-Whitney test, *p* < 0.05; **Fig. 3b** inset) than positions <150 nucleotides into the read. Our data therefore show that false positive iSNVs are not evenly distributed across the Illumina sequencing reads, and that the profiles are consistent with published Illumina error rates [16]. We found that the occurrence and iSNV frequency of false positives, decreased during the last 50 nucleotides of the Illumina reads (positions 200-250, **Fig. 3a-b**), which is due to overlapping reads during paired-end sequencing of 400 bp amplicons (data not shown).

**Figure 3.**
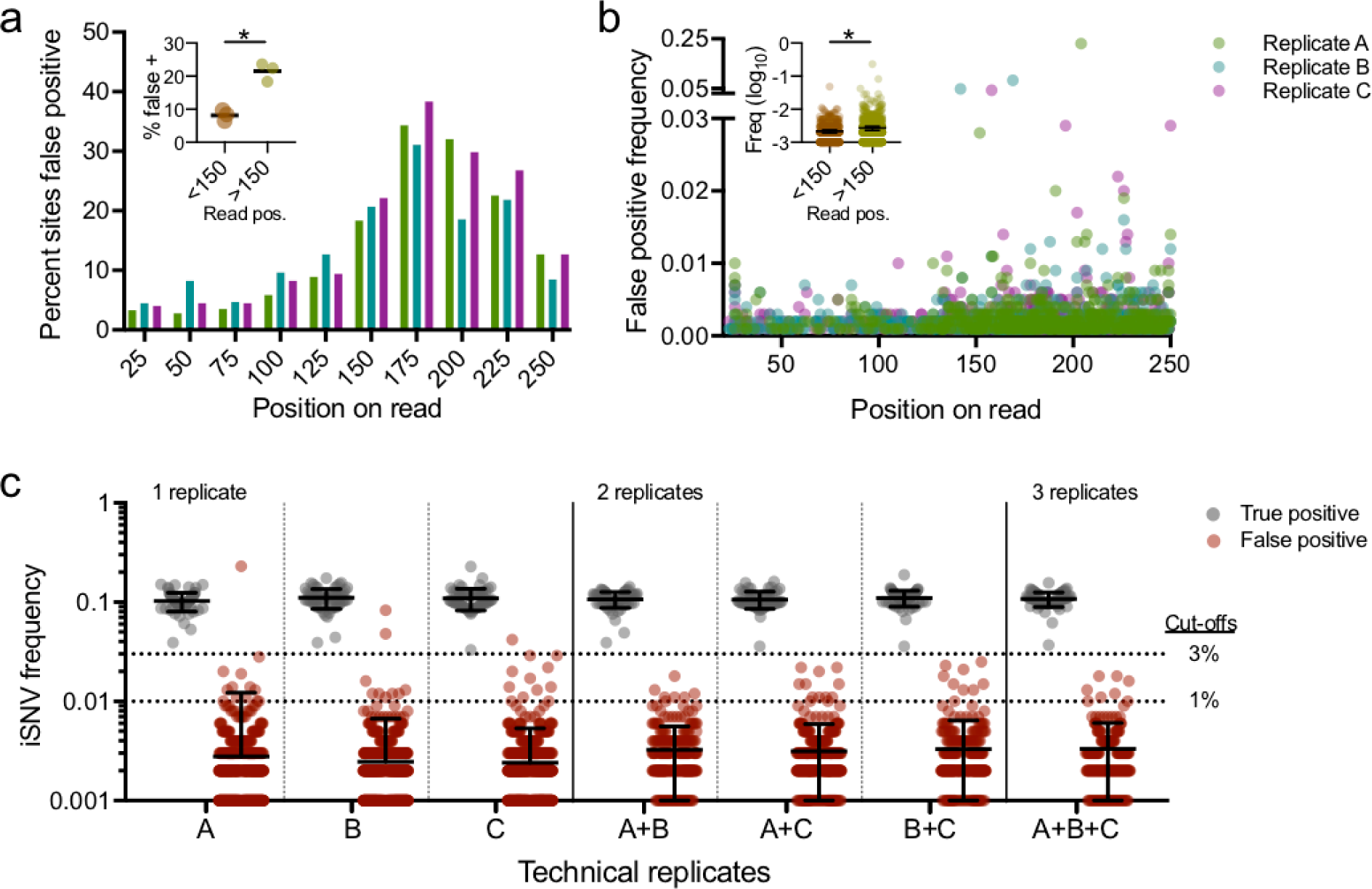
False positive intrahost variants caused by sequencing errors can be removed by technical replicates and frequency cut-offs. We sequenced mixed Zika virus population containing 10% of virus #2 in triplicate, limited our analysis to the regions only covered by perfect PCR primer matches, and removed sites with intrahost single nucleotide variant (iSNV) detected at >1% frequency in either of the Zika virus isolates. This left us with 61 true positive (10% frequency) iSNV sites and 3940 sites not expected to be variable to investigate false positives (>0.1% frequency). (**a**) The locations of false positives on the sequencing read position were mapped and shown as the distribution within 25 nt bins by percent of sites with false positive iSNV calls. Each color represents data from an independent replicate. Inset: Read positions >150 nt had a significantly higher false positive rate than positions < 150 nt (*, Wilcoxon test, *p* < 0.05). (**b**) The iSNV frequencies from each false positive were also plotted by position on the sequencing read. Each color represents data from an independent replicate. Inset: False positive iSNV frequencies were significantly higher at read positions > 150 nt than < 150 nt (*, Mann-Whitney test, *p* < 0.05). (**c**) True and false iSNVs were plotted by frequency for each individual replicate (A, B, and C) and combined as technical duplicates and triplicates showing the mean frequencies of iSNVs only found in all replicates. Data shown as means and standard deviations. The line indicates the proposed cut-off at 3% based on removing false positives from the replicate data while still in the range of high accuracy (**Fig. 1b**).

Knowing the general distribution of false positive iSNVs, we sought to remove them post sequencing. Based on previous investigations [12,35,36], we proposed to remove false positive iSNVs by (***1***) amplifying and sequencing each sample as technical replicates (at least twice), and (***2***) only calling iSNVs detected in all replicates. Using our Mix10% virus population, we analyzed each replicate in isolation or in combination, and calculated the mean iSNV frequencies (**Fig. 3c**). From individually sequenced replicates, we found 1-2 false positive iSNVs per sample were within the frequency distribution of our true iSNVs (**Fig. 3c**, panel “1 replicate”), demonstrating that a simple frequency cut-off will either leave false positive or remove true positive iSNVs. When considering replicates in combination, however, we found that the percent of sites with a false positive iSNV call (above 0.1%) dropped from ~16% (**Fig. 3c** “1 replicate”) to ~9% (**Fig. 3c** “2 replicates”). More importantly, we found that all of the false iSNVs that passed the duplicate filter had frequencies below the 3% limit of accurate iSNV measurements (**Fig. 1b**). This allowed us to use a secondary frequency cut-off (3%) to remove the remainder of the false positives, while maintaining all of the true (10%) iSNVs. We found that the addition of a third technical replicate only resulted in a moderate reduction of sites with false iSNVs above 0.1% (9% to 6%) and did not help us to decrease the frequency cut-off filter (**Fig. 3c** “3 replicates”). We conclude that PrimalSeq can be used for accurate iSNV detection above 3% when using at least two technical replicates.

### The accuracy of PrimalSeq is comparable to metagenomic sequencing

The gold standard for virus sequencing is untargeted metagenomics - sampling all RNA or DNA present in a sample [34]. Compared to amplicon-based sequencing, metagenomic sequencing use random priming and do not select for specific RNA sequences and is thus less biased. In addition, for virus sequencing in resource-limited conditions, the Oxford Nanopore MinION is gaining popularity due to its portability and low instrument cost [19,28,37], which is enabling real-time outbreak tracking [38]. To compare iSNV calling accuracy across platforms and methods, we evaluated iSNV measurements using either: (***1***) metagenomic Illumina sequencing, (***2***) PrimalSeq with Illumina sequencing, and (***3***) PrimalSeq with Oxford Nanopore sequencing. We found that PCR amplification leads to more variable iSNV frequency measurements, but that the overall accuracy of PrimalSeq is comparable to metagenomic sequencing (**Fig. 4**). PrimalSeq using Nanopore sequencing can be used to detect iSNVs, but, as expected, high error rates [39,40] makes it difficult to differentiate between true and false iSNVs (**Figs. 4 & 5**).

**Figure 4.**
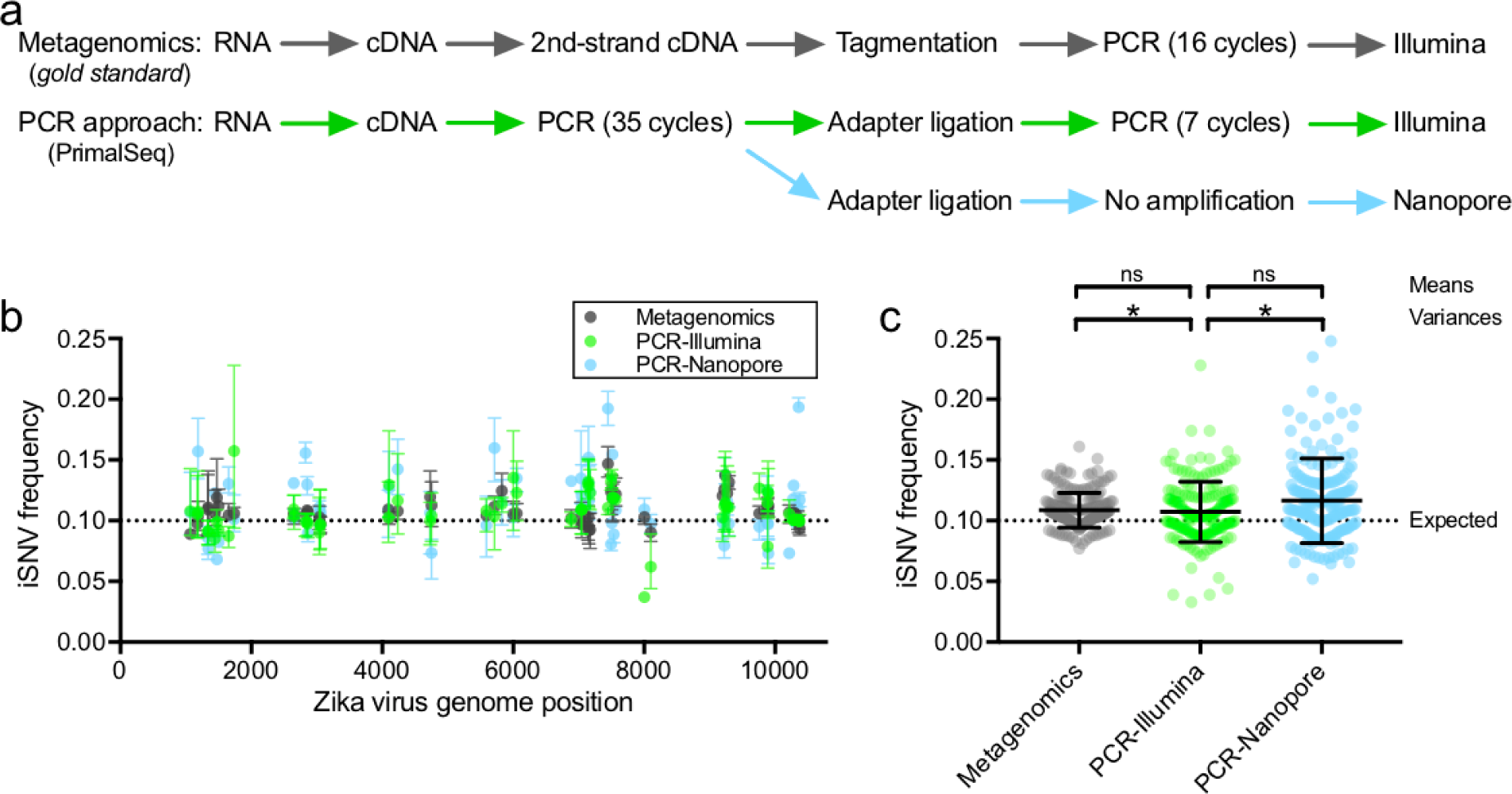
PCR amplification prior to sequencing leads to similar overall measurements of genetic diversity. (**a**) We compared our PrimalSeq that enriches for specific virus sequences to the current ‘gold standard’ for measuring intrahost genetic diversity, metagenomics; and we compared sequencing the amplicons using the Illumina and Oxford Nanopore platforms. The schematic outlines the general workflow for all approaches. (**b**) We sequenced our mixed Zika virus population (1000 virus RNA copies) containing 10% virus # 2 (Expected) in triplicate using both approaches and platforms to compare the accuracy of measuring known intrahost single nucleotide variants (iSNVs). We only analyzed regions of the Zika virus #1 and #2 genomes (Fig. 1a) that were perfect matches to the PCR primer sequences, leaving 61 iSNV sites. Data shown as mean and range of triplicate tests. (**c**) We combined the frequency measurements for each iSNV site and replicate (*n* = 183) to compare the accuracy between the two approaches and platforms. Data shown as means and standard deviations. The mean frequencies were not significantly different (ns, Welch’s t test, *p* > 0.05), but the variances were not equal (*, Levene’s test, *p* < 0.05).

**Figure 5.**
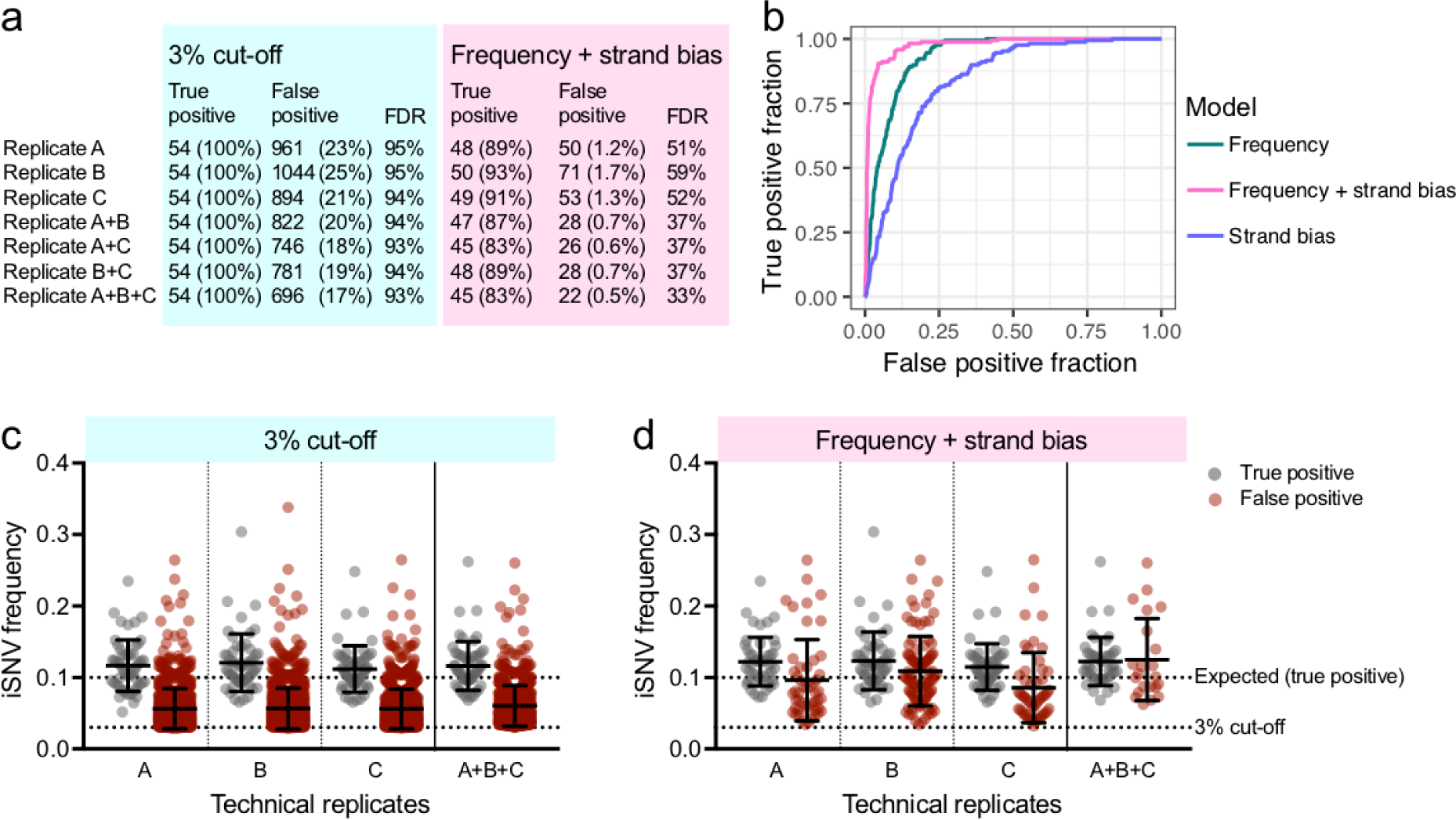
High false discovery rates of intrahost variants using Nanopore sequencing. (**a**) iSNV false discovery rates (FDR) from Oxford Nanopore sequencing data. We analyzed 54 true positive and 4173 true negative sites, and determined the proportion of true and false positive iSNV calls from datasets containing 1, 2, or 3 technical replicates using either a 3% frequency cut-off or a logistic regression of iSNV frequency and strand bias. (**b**) A receiver operator characteristic (ROC) curve showing a logistic regression model that incorporates allele frequency and strand bias as features and the presence or absence of a iSNV as the response variable. The model was trained and tested using a ten fold cross validation scheme. The model was performed using a frequency and strand bias threshold alone, and combining the two features. Post-filter iSNV frequencies of true and false positive calls using (**c**) a 3% cut-off or (**d**) a a logistic regression of iSNV frequency and strand bias. Data shown as the means and 95% confidence intervals.

To compare PrimalSeq and metagenomic sequencing approaches, as well as platforms (Illumina or Nanopore), for iSNV measurement accuracy, we generated triplicate sequencing libraries from our Mix10% virus population (1000 RNA copies). The triplicate amplicon libraries were sequenced using both Illumina MiSeq and Oxford Nanopore MinION platforms, while the metagenomics libraries were sequenced using the MiSeq (**Fig. 4a**). We found that compared to metagenomic sequencing, mean iSNV frequencies generated from PrimalSeq were not significantly different when using the Illumina platform (Welch’s t test, *p* > 0.05; **Fig. 4b-c**). The variances of individual iSNV frequencies, however, were significantly higher using PCR amplification (Levene’s test for variance, *p* < 0.05; **Fig. 4c**). Compared to Illumina sequencing, the mean iSNV frequencies measured using the Oxford Nanopore MinION were not significantly different (Welch’s t test, *p* > 0.05; **Fig. 4b-c**), however, variances of individual iSNV frequencies were significantly higher using Nanopore (Levene’s test for variance, *p* < 0.05; **Fig. 4c**). These findings show that while PCR amplification lead to more variable results for individual iSNVs, the overall measured diversity is comparable to metagenomic sequencing. Furthermore, we show that iSNV frequencies become even more variable when sequencing using the Oxford Nanopore platform, which is consistent with its higher error rate [39,40].

### High intrahost variant false discovery rate using Oxford Nanopore sequencing

To further explore Oxford Nanopore sequencing for measuring intrahost virus diversity, we examined if we could (***1***) differentiate between mixed genotypes within a virus population and (***2***) computationally remove false positive iSNVs. Though the mean iSNV frequencies measured from Nanopore were not significantly different from Illumina (**Fig. 4**) and we could assign reads to the correct haplotype (*i.e.*, virus #1 or #2), we found it difficult to differentiate between true and false positive iSNVs (**Fig. 5**). For this evaluation, we used our Nanopore data generated from the Mix10% Zika virus population (**Fig. 4a**).

First, using a reference database containing virus #1 and #2, plus two divergent Zika viruses (French Polynesia, 2007; Uganda, 1947), we determined if we could differentiate between virus haplotypes in a mixed population. We found that 92.38% of the aligned reads mapped to virus #1 and 7.35% mapped to virus #2, the roughly expected 90%:10% proportions (0.27% of reads mapped erroneously to haplotypes not present in the mixture). Overall, the results indicate that nanopore sequencing reads are useful for identifying unique haplotypes within a mixture - as might be expected for co-infections [41] - despite a high error rate.

To attempt to differentiate between true and false positive iSNVs, we limited our analysis to regions only covered by perfect primer matches and analyzed 54 true positive and 4173 true negative sites, as we did above for the Illumina data (**Fig. 3**). We filtered the sequencing data using technical replicates and a 3% frequency cutoff, which we demonstrated above could be used to remove false iSNV calls in our Illumina data (**Fig. 3c**). Using these filters, we found that >17% of the 4173 invariant sites had false iSNV calls in the Nanopore data, even when including all three replicates (**Fig. 5a**). This is because the majority of the false positive iSNVs had measured frequencies as high, or higher, than the 10% true positives (**Fig. 5c**), leading to a >93% false discovery rate (**Fig. 5a**). To investigate if the false discovery rate could be reduced using additional data within the sequencing reads, we trained a logistic regression model incorporating iSNV frequency and strand bias [42] as features, and the presence or absence of known iSNVs as the response variable (**Fig. 5b**). Based on this analysis, we found that using a frequency and strand bias filter resulted in a higher true or false iSNV discriminatory power, as shown by its greater area under the curve, than the two features independently (**Fig. 5b**). Using this filter for individual replicates, we were able to reduce the number of false positive iSNVs from ~900-1000 (3% cut-off) to ~50-70 (frequency + strand bias) and the false discovery rate from ~95% to ~55% (**Fig. 5a**). By including replicate sequencing (either 2 or 3), we could further reduce the false discovery rate to <40% (**Fig. 5a**). Despite this significant reduction, the remaining false positive iSNVs still had high frequencies in our dataset (~5%-25%, **Fig. 5d**). These findings show that estimating intrahost virus genetic diversity using the Oxford Nanopore platform will require additional technological and computational innovations for anything other than simple scenarios of co-infections with diverse virus haplotypes.

### Accurate intrahost variant calling using iVar

Using the above validation experiments, we generated a comprehensive experimental protocol (**Additional file 2**) and an open source computational tool, iVar (**i**ntrahost **v**ariant **a**nalysis of **r**eplicates; github.com/andersen-lab/ivar) to accurately measure iSNVs (**Fig. 6**). Our framework is compatible with any PCR-based sequencing approaches but was specifically designed for use with PrimalSeq on the Illumina platform. For the experimental protocol, we added the following recommendations to PrimalSeq [17]: (***1***) start with at least 1000 RNA copies of the virus (**Fig. 1c**), (***2***) prepare each sample as technical duplicates (**Fig. 3c, Fig. 4c**), and (***3***) sequence to a depth of at least 400× (**Fig. 1d**).

**Figure 6.**
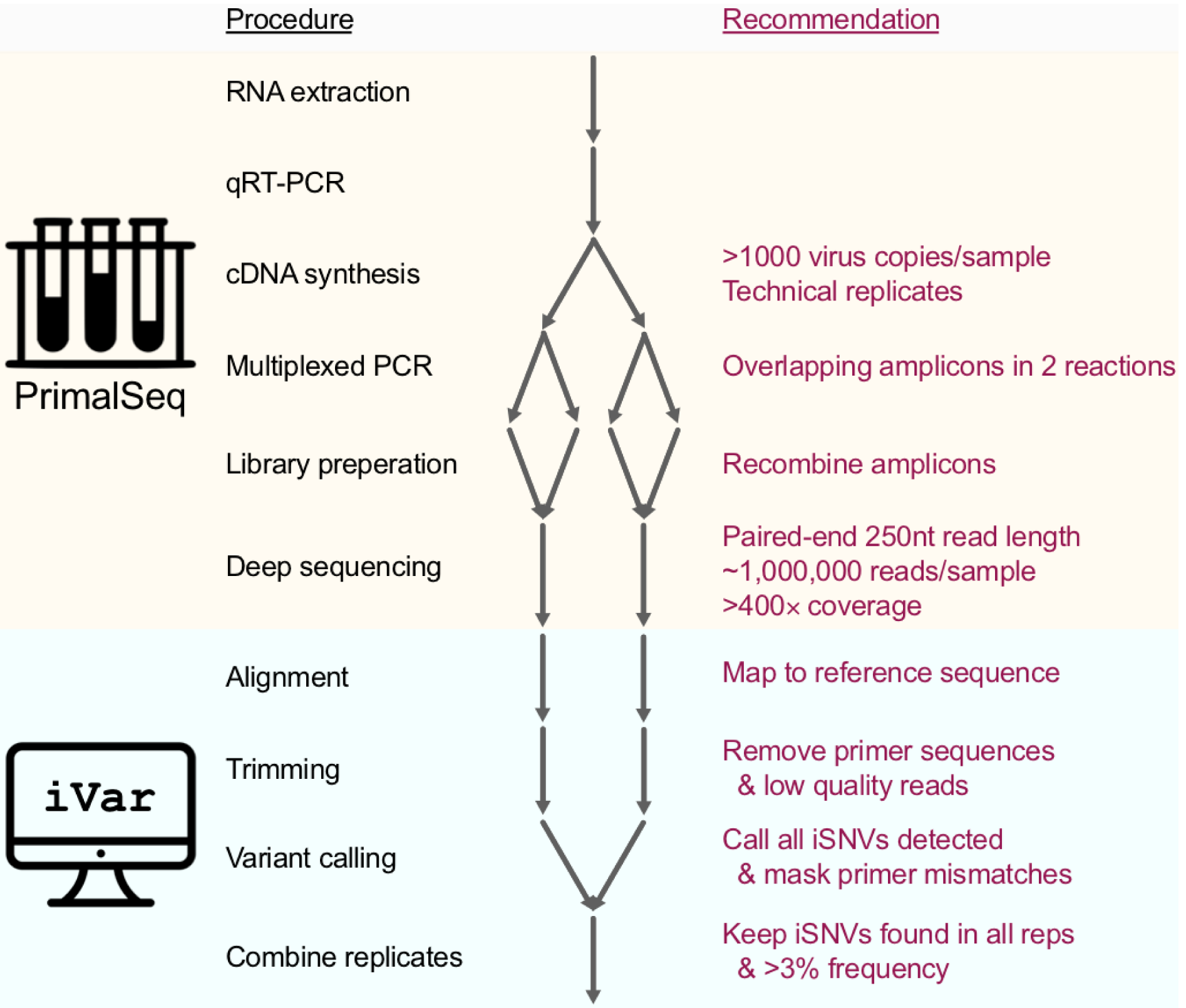
Experimental and computational workflow for measuring intrahost virus diversity using PrimalSeq.

Our computational package contains functions broadly useful for viral amplicon-based sequencing. Additional tools for metagenomic sequencing are actively being incorporated into iVar. While each of these functions can be accomplished using existing tools, iVar contains an intersection of functionality from multiple tools that are required to call iSNVs and consensus sequences from viral sequencing data across multiple replicates. We implemented the following functions in iVar: (***1***) trimming of primers and low-quality bases, (***2***) consensus calling, (***3***) variant calling - both iSNVs and insertions/deletions, and (***4***) identifying mismatches to primer sequences and excluding the corresponding reads from alignment files. We created two pipelines in iVar to call iSNVs from samples prepared using an amplicon-based library preparation method, with or without a known reference sequence (*e.g.*, experimental or field-derived), and with or without technical replicates (with no limit on the number of technical replicates). From our empirically-derived data, we recommend the following: (***1***) only call iSNVs detected in both replicates greater than 3% frequency (**Fig. 1a**, **Fig. 3c**) and (***2***) remove reads from amplicons with mismatched primers to normalize comparisons of intrahost population genetics (**Fig. 2**).

We used existing functions in VarScan2 [43], MAFFT [44], Geneious [45], Trimmomatic [46] and cutadapt [47] to test the accuracy of the trimming, consensus sequence generation, and variant calling functions in iVar. We found that iVar performed as well as, or better than, each of the tools we benchmarked it against (**Fig. S1-S3**). We used two simulated datasets and two clinical Zika virus samples sequenced using PrimalSeq to validate iSNV calling, and found an almost perfect correlation between iVar and VarScan2 (Spearman’s ρ=1; **Fig. S1**). We also found zero nucleotide differences in the consensus sequences called using iVar and Geneious at all four thresholds (0%, 25%, 50%, and 90%) and across the four datasets (**Fig. S2**). We found that iVar was better at trimming primer sequences than cutadapt (**Fig. S3**). This is because iVar uses primer positions specified in a BED file to soft clip the primer regions after alignment whereas cutadapt trims sequencing reads by comparing the primer nucleotide sequence with the nucleotides at the 5’ end of each read, before alignment. As a result, iVar was able to trim sequencing reads which might not start or end exactly at the beginning of the primer sequence (**Fig. S3**). Since cutadapt uses the actual primer sequences, which are assumed to be anchored at the 5’ or 3’ end to do the trimming, it misses these cases. We trimmed the length of the longest primer sequence (22 bp) from the 5’ end of all the sequenced reads using the “HEADCROP” option in Trimmomatic. This approach, however, is crude and will result in a loss of 22 bp form the 5’ end of all sequenced reads. Thus, iVar contains functionality that is fine tuned to perform iSNV calling from data generated using amplicon-based protocols.

### PrimalSeq and iVar can be used to measure intrahost virus genetic diversity from primary samples

There are many sequencing options available to measure intrahost virus diversity from cell culture stocks (*e.g.*, [36]) or infected animals with high titers (*e.g.*, [34]). These approaches, however, often do not generate sufficient data when there is high host background RNA, or low virus copies, as is the case for many viruses, including Zika, during human and primate infections [18–21]. Using our validated PrimalSeq and iVar framework (**Fig. 6**), and by using Primal Scheme to create a new multiplexed primer set to amplify West Nile virus, we thoroughly evaluated our approaches to measure intrahost diversity from 36 Zika virus and West Nile virus populations (**Fig. 7**). These samples came from a variety of laboratory and field-derived sample types and were each amplified, sequenced, and analyzed in duplicate (**Table S2**, **Fig. 7**). We omitted (*i.e.*, “masked”) from comparative genomic analysis regions of the virus genomes with iSNV mismatches in the primer binding regions (~2 per sample) or with sequencing coverage depth <400× (**Table S2**, **Fig. 7**, grey bands).

**Figure 7.**
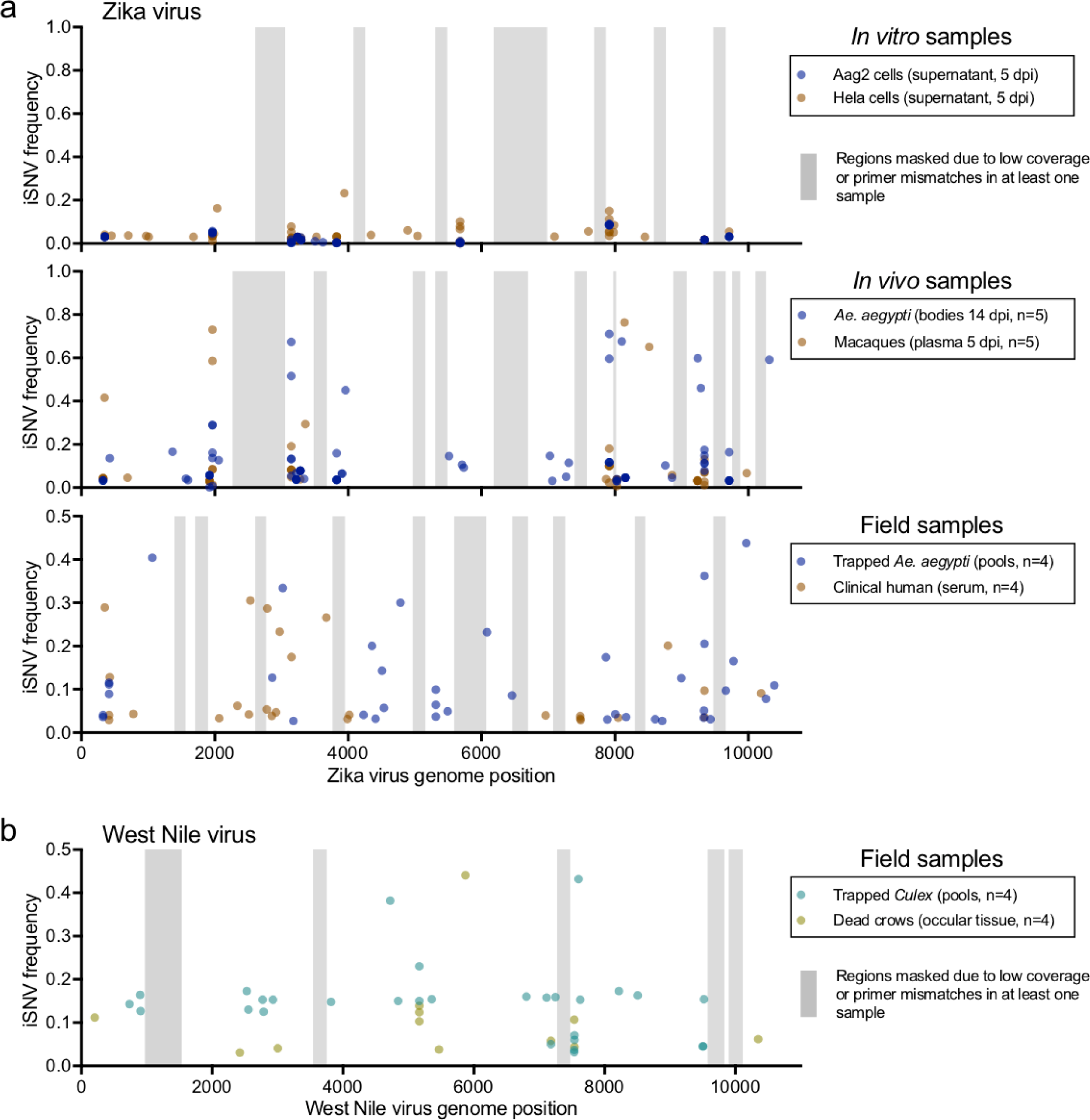
PrimalSeq can be used to measure intrahost variants from a variety of sample types. (**a**) We sequenced technical duplicates of Zika virus populations (1000 virus RNA copies each) to identify intrahost single nucleotide variants (iSNVs) >3% within each sample. *In vitro* and *in vivo* samples were generated using Zika virus strain PRVABC59 (isolated from Puerto Rico, 2015) during infection of *Ae. aegypti* Aag2 cells (derived from embryos), human HeLa cells (derived from cervical epithelial cells), *Ae. aegypti* mosquitoes (orally infected), and Indian origin rhesus macaques (subcutaneously infected). For the *in vitro* and *in vivo* samples, where the reference population sequence is known, the iSNV frequencies were calculated by change in frequency from pre-to post-infection. Field Zika virus samples from pooled *Ae. aegypti* and human clinical samples were collected from Florida during the 2016 Zika virus outbreak. (**b**) *Culex* mosquitoes and dead American crows were collected from San Diego County, CA, during 2015 to sequence West Nile virus from field samples (10,000 virus RNA copies each). The iSNV frequencies from the field samples are the minor allele frequencies (maximum frequency = 0.5) because the reference virus sequence was not known. For both **a** and **b**, analysis was limited to regions of the genome with >400x coverage depth in the protein coding sequence and we masked amplicons with primer mismatches from our analysis (grey regions) for direct comparisons of intrahost genetic diversity.

To demonstrate the types of analyses that can be performed with PrimalSeq and iVar, we compared the mosquito- and vertebrate-derived virus samples using several measures of intrahost diversity and selection (**Fig. 8**). We measured genetic richness (the number of iSNV sites; **Fig. 8a**), complexity (uncertainty associated with randomly sampling an allele; **Fig. 8b**), distance (the sum of all iSNV frequencies; **Fig. 8c**), and selection (the proportion of nonsynonymous iSNVs; **Fig. 8d**). We did not analyze masked regions, so that only high confidence regions of the genome from all samples within the experiment were compared (**Fig. 7**). We found that Zika virus genetic diversity (all measures) and selection was significantly higher from populations derived from primate (Hela) cells than *Ae. aegypti* (Aag2) cells (**Fig. 8**). *In vivo*, however, our findings were reversed. Zika virus genetic richness, complexity, and selection were significantly higher in *Ae. aegypti* bodies than primate (rhesus macaque) plasma (**Fig. 8**). Furthermore, we found that the distribution of iSNV frequencies of the Zika virus populations was similar across different *in vivo* infections (**Fig. 8e**). This finding indicates that the increased Zika virus diversity in mosquitoes was driven by more 3-20% iSNVs that were also common in macaques, and not by a few additional high frequency iSNVs. We found that from both Zika and West Nile virus field samples, genetic diversity and selection were not significantly different between virus populations isolated from their mosquito vectors (*Ae. aegypti* or *Culex*) or vertebrate hosts (humans or birds), though the mosquito samples were more variable (**Fig. 8**). Overall, our data suggest that virus diversity is highly dependent on the experimental and biological systems, and future uses of PrimalSeq will help identify the mechanisms underlying these evolutionary differences (**Fig. 8f**).

**Figure 8.**
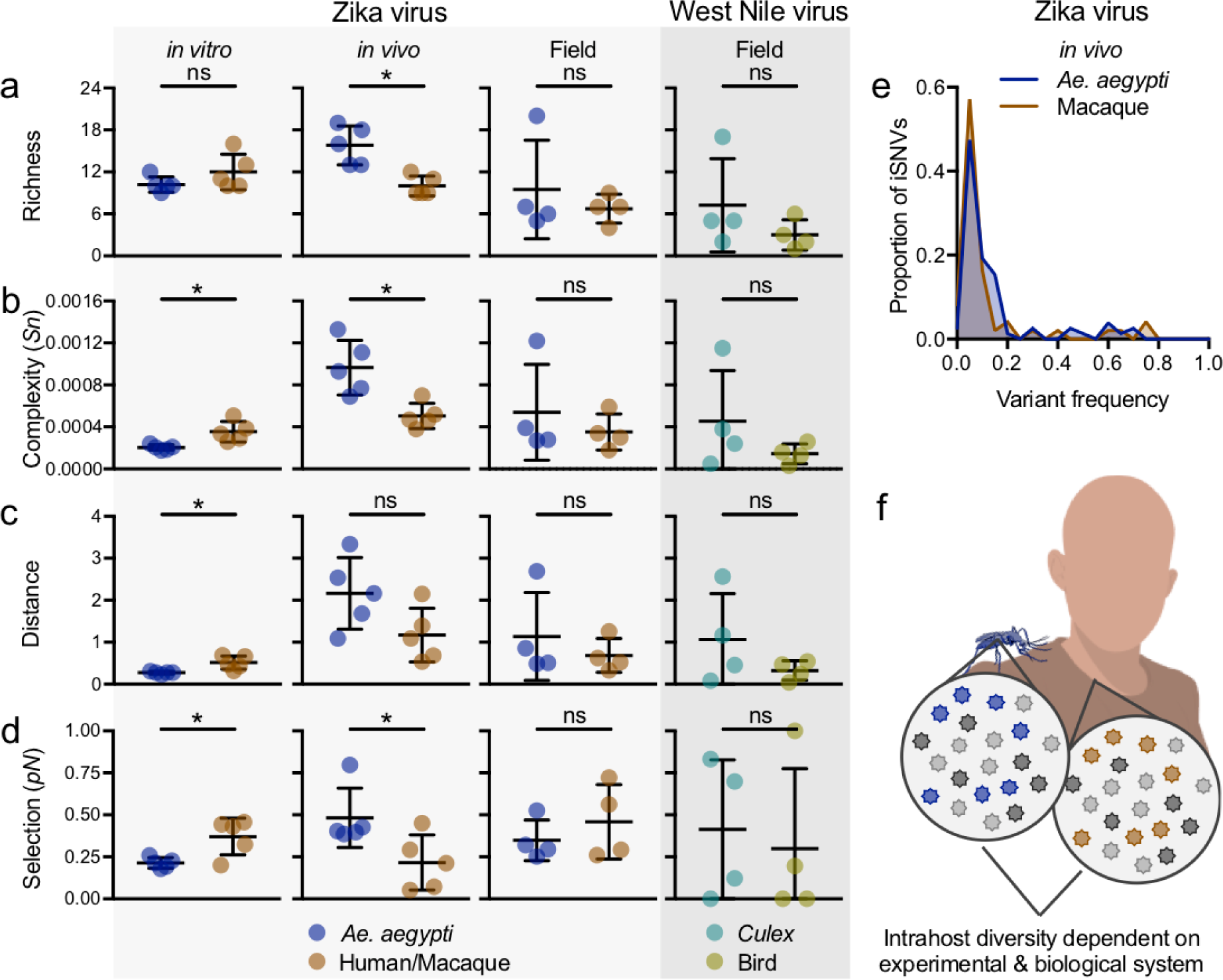
Intrahost virus genetic diversity is dependent on the experimental and biological system. Variants called from Zika and West Nile virus populations derived from *in vitro*, *in vivo*, and field studies (Fig. 6) were used to compare intrahost virus diversity and selection from mosquito vectors (*Ae. aegypti* and *Culex* species) and vertebrate hosts (primates or birds). We compared (**a**) richness (the number of intrahost single nucleotide variant [iSNV] sites; Fig. 7a), (**b**) complexity (uncertainty associated with randomly sampling an allele, measured by Shannon entropy [*Sn*]), (**c**) distance (the sum of all iSNV frequencies), and (**d**) selection (the proportion of nonsynonymous iSNVs [*pN*]). The mosquito and vertebrate-derived populations were compared using unpaired t-tests (ns, not significant; *, *p* < 0.05). Data shown as mean and standard deviation. (**e**) The proportion of Zika virus iSNVs detected in the *Ae. aegypti* and rhesus macaque *in vivo* samples were distributed by frequency. Bin width is 0.05. (**f**) Our combined data suggests that intrahost virus diversity is dependent upon the experimental system (*i.e.*, *in vitro*, *in vivo*, or field samples).

## Discussion

Understanding how RNA viruses evolve within hosts can help lead to breakthroughs in medicine and biology. Rapid developments in sequencing technologies are facilitating more research into these areas, yet, usage of unvalidated tools and systematic biases can dramatically limit the utility of the results [12–16]. To address these concerns, we developed PrimalSeq [17] and validated it across different platforms and sample types. In our development of iVar we show that PrimalSeq can be used to accurately measure intrahost virus diversity from different viruses and samples with amplicon-based sequencing. Based on our experimental validations, we provide a detailed laboratory protocol (**Additional file 2**), and suggest the following best practices for measuring intrahost virus diversity when using amplicon-based sequencing approaches:

1. Start with at least 1000 RNA copies of the virus for the initial cDNA synthesis step.
2. Prepare the RNA from virus population for sequencing in duplicate.
3. Sequence each library to a depth of at least 400× at each genome position using the Illumina platform.
4. Only call iSNVs greater than 3% frequency that are detected in both replicates (a lower frequency may be achievable with higher RNA quantities).
5. For multi-sample comparisons of genetic diversity, omit genome regions amplified with primers that contain iSNVs within the binding sites.

Several factors can alter the accuracy of measuring intrahost virus diversity. In particular, we found that input virus concentrations, sequencing coverage depths, and primer mismatches can have profound effects on iSNV estimations. Using the recommendations above, however, we could consistently and accurately detect iSNVs at 3% frequency and higher. We predict that the lower limit of iSNV detection can be improved with a higher effective sampling depth (*i.e.*, more input virus and deeper coverage) [13].

Given no primer mismatches to the virus sequences, we found that measures of iSNVs frequencies from PrimalSeq were nearly as accurate as an untargeted metagenomics approach [34]. Because iVar remove the primer sequences from downstream analysis and use overlapping amplicons, frequency measures of iSNVs within the primer regions themselves are not skewed. Instead, iSNVs within primer regions can alter the measured frequencies of other iSNVs within that particular amplicon. In these cases, results should be interpreted with caution, and we incorporated a step in iVar to mask out such regions for comparative analyses. It is plausible that using primers with degenerate nucleotides at mismatched iSNV positions could help alleviating this bias [33].

False iSNV calls significantly influence measurements of intrahost virus diversity [12]. We found that the positive association of false positive iSNVs with sequencing read lengths better fit the profiles of Illumina sequencing errors, rather than PCR errors [16,48]. In fact, we estimate that the Illumina MiSeq error rate (~0.9% [16]) is ~60× greater than the error rates during PCR in our approach (~0.02% [49]). Therefore, PrimalSeq likely does not add significantly more error, and by extension false iSNVs, than what was already inherent to the Illumina sequencing platform. Indeed, we found that PrimalSeq was comparable to PCR-free metagenomic sequencing in estimating intrahost virus diversity.

The ease and portability of Oxford Nanopore technologies, particularly the MinION, are revolutionizing the way we sequence viruses, including its use in near real-time outbreak tracking [19,28,37]. Our data indicate, however, that the Nanopore platform is not yet adequate for detection of minor alleles and measures of intrahost diversity. While it may provide value in tracking frequency changes of known iSNVs over time, we found that the high error rates (10-15% [39,40]) makes it difficult to differentiate between true and false iSNV calls. We found that stringent post-filtering, such as combining iSNV frequencies and strand bias across comparing replicate samples, significantly reduce false positive iSNV calls, but there is still a high false discovery rate. Effectively using Nanopore sequencing for intrahost virus diversity measurements will require higher sequencing accuracy and base calling, exploitation of co-occurring variants (*i.e.* haplotyping) [50], or utilization of different molecular approaches, including the 1D^2^ method (where template and complementary strands of each fragment are sequenced) [40], tandem repeat consensus techniques [36,51] or unique molecular identifiers [52].

For viruses that utilize multiple hosts, like mosquito-borne viruses, being able to compare results from many samples types is critically important. A lack of standardization, however, means that the field does not yet have a consensus to whether the mosquito vector or the vertebrate host contributes the most to virus genetic diversity [53–62]. The development of PrimalSeq and iVar allows for such measurements to be performed across diverse environments, sample types, and experimental designs. Using PrimalSeq, for example, we found that *in vitro* Zika virus diversity was significantly greater in human cells, when compared to mosquito cells. However, we found that these results were reversed during *in vivo* studies. Furthermore, we did not detect significant differences in field-collected mosquito and vertebrate samples for both Zika and West Nile virus. A caveat for the field samples, however, is that we do not know the reference sequence and cannot account for consensus-changing mutations introduced through intrahost bottlenecks and genetic drift [63–65]. Our incongruent results among experimental designs help to explain why there is still debate about the relative impact of vectors and hosts on virus evolution, and further use of PrimalSeq will help to resolve these issues.

## Conclusions

We demonstrate that PrimalSeq can accurately measure intrahost virus genetic diversity if properly validated. We benchmarked our highly multiplexed and streamlined amplicon-based sequencing method using a series of experiments with mixed virus populations, developed an all-inclusive computational analysis tool (iVar), and showcase its utility by measuring intrahost virus diversity from cells, mosquitoes, primates, birds, and humans. Furthermore, using our free online primer designer, Primal Scheme (primal.zibraproject.org) [17], PrimalSeq can be modified for use with a wide range of viruses. Overall, our detailed laboratory and computational approaches presented here can reveal important insights about intrahost virus evolution directly from clinical or experimental samples in a way that is cheap, accurate, and scalable.

## Methods

### Ethics approval and consent to participate

Research on human subjects was conducted in compliance with existing regulations relating to the protection of human subjects and was evaluated and approved by the Institutional Review Board/Ethics Review Committee at The Scripps Research Institute. Clinical samples were obtained from the Florida Department of Health (DOH) and Antibody Systems Inc. Samples collected in Florida were collected under a waiver of consent granted by the Florida DOH Human Research Protection Program. The work received a non-human subjects research designation (category 4 exemption) by the Florida DOH since this research was performed with leftover clinical diagnostic samples involving no more than minimal risk. All samples were de-identified prior to receipt by the study investigators. Research involving Indian origin rhesus macaques was conducted at the California National Primate Research Center, and experimental infections of mice upon which *Ae. aegypti* mosquitoes fed were performed at the University of California, Davis, School of Veterinary Medicine. Both institutes are fully accredited by the Association for the Assessment and Accreditation of Laboratory Animal Care International. Animals were cared for in accordance with the National Research Council Guide for the Care and Use of Laboratory Animals and the Animal Welfare Act. Animal experiments were approved by the Institutional Animal Care and Use Committee of UC Davis (protocols #19211 and #19695 for rhesus macaques, protocol #19404 for mice). All macaques samples used in this study were from approved studies [66]; and none were generated specifically for this work.

### Mixed virus populations

Zika virus RNA from isolates PRVABC59 (Puerto Rico 2015, Genbank KX087101, ‘virus #1’) and FSS13025 (Cambodia 2010, Genbank KU955593, ‘virus #2’) were quantified by qRT-PCR (as previously described [26]). The consensus sequences from PRVABC59 and KX087101 were determined using untargeted metagenomics (see below) and a strict >99% majority nucleotide threshold at each site. Sites that were mixed (*i.e.*, containing an iSNV >1% frequency) were not used to evaluate iSNVs at known frequencies (**Fig. 1**). Using quantified virus RNA copies, the two viruses were mixed to achieve the desired total RNA copies (½ required amount because 2 μL of RNA was used for cDNA) and ratios of PRVABC59:FSS13025. Metagenomic sequencing of a 10:1 mixed virus population (*i.e.*, 10% FSS13025) was used to verify our mixing approach (**Fig. 4**). Each mixed virus population was sequenced in triplicate using the metagenomic and amplicon approaches described below.

### Laboratory-infected cells, mosquitoes, and primates

Zika virus was collected from *in vitro* and *in vivo* experiments to compare intrahost diversity between mosquitoes (*Ae. aegypti*) and primates (humans and macaques, **Table S2**). All *in vitro* and *in vivo* experiments were conducted using Zika virus isolate PRVABC59 (Puerto Rico, 2015, KX087101). All Zika virus RNA was quantified by qRT-PCR, as described [26].

Aag2 (derived from *Ae. aegypti* embryos [67]) and HeLa (derived from human cervical epithelial cells, ATCC CCL-2) cells were infected using a multiplicity of infection of 0.01 and supernatant was harvested 5 days post infection. Both cell lines were maintained using Minimal Essential Medium (Sigma-Aldrich) supplemented with 10% (v/v) fetal bovine serum, L glutamine, sodium bicarbonate, and antibiotics (penicillin and streptomycin). Aag2 and HeLa cells were incubated with 5% CO_2_ at 27 ℃ and 37 ℃, respectively.

*Ae. aegypti* mosquitoes were infected with Zika virus as previously described [68]. In brief, colonized mosquitoes originating from Los Angeles, California in 2016 feed on viremic mice inoculated with 5 log_10_ Vero plaque forming units of Zika virus (PRVABC59). At 14 days post infection, individual mosquitoes were collected and homogenized. Viral RNA was extracted from 50 μL of mosquito homogenate using the using the MagMax Viral RNA Extraction Kit and eluted 50 µL of elution buffer (Buffer EB, Qiagen). Indian origin rhesus macaques (*Macaca mulatta*) were inoculated subcutaneously with 3 log_10_ Vero plaque forming units of Zika virus (PRVABC59) and plasma was collected 5 days post infection, as described [66,69]. RNA was extracted from at least 300 μL of rhesus macaque plasma using the MagMax Viral RNA Extraction Kit and was eluted in 60 μL of elution buffer. RNA extracts from laboratory infected mosquitoes and macaque plasma used for this study had been thawed previously at least one time.

### Field-collected mosquitoes and clinical samples

Clinical and entomological samples were collected during the 2016 Florida Zika virus outbreak [26] to compare intrahost Zika virus diversity between naturally infected humans and mosquitoes (**Table S2**). Human clinical samples were obtained for diagnostic and surveillance purposes and excess human sera were used for this study. RNA was extracted using the RNAeasy kit (Qiagen) and eluted into 50-100 µL using the supplied elution buffer. Entomological samples were collected by the Miami-Dade Mosquito Control for surveillance of Zika virus activity. *Ae. aegypti* mosquitoes were collected using BG-Sentinel mosquito traps (Biogents AG) and sorted into pools of up to 50 females per trap. The pooled mosquitoes were stored in RNAlater (Invitrogen), RNA was extracted using the RNAeasy kit (Qiagen), and Zika virus RNA was quantified by qRT-PCR [26]. RNA from Zika virus positive pools used in this study contained 13-39 individual mosquitoes; however, considering that ~1 in 1600 were infected [26], it is highly unlikely that any pool contained >1 infected mosquito.

*Culex quinquefasciatus* mosquitoes (up to 50 per trap) and dead American crows were collected by the San Diego County Vector Control Program during 2015. RNA was extracted using the RNAeasy kit (Qiagen) and screened for the presence of West Nile virus RNA using standard RT-qPCR.

### PCR amplification of the virus genomes

Virus RNA (2 μL) was reverse transcribed into cDNA using Invitrogen SuperScript IV VILO (20 µL reactions). Virus cDNA (2 μL) was amplified in 35 × ~400 bp fragments from two multiplexed PCR reactions using Q5 DNA High-fidelity Polymerase (New England Biolabs) using the conditions previously described [17]. For the data shown in **Fig. 1**, the mixed Zika virus populations were amplified in one multiplexed reaction containing primer sets 5, 24, and 33. A detailed protocol can be found in Additional file 2 and the Zika and West Nile virus primers can be found in **Additional file 1** (**Table S3** and **Table S4**, respectively).

### Amplicon-based Illumina sequencing

A detailed protocol for our amplicon-based sequencing methods can be found in **Additional file 2**. Protocol updates will be released online at http://grubaughlab.com/open-science/amplicon-sequencing/ and https://andersen-lab.com/secrets/protocols/. Virus amplicons from the two multiplex PCR reactions (above section) were purified using Agencourt AMPure XP beads (Beckman Coulter) and combined (25 ng each) prior to library preparation. The libraries were prepared using the Kapa Hyper prep kit (Kapa Biosystems, following the vendor’s protocols but with ¼ of the recommended reagents) and NEXTflex Dual-Indexed DNA Barcodes (BIOO Scientific, diluted to 250 nM). Agencourt AMPure XP beads (Beckman Coulter) were used for all purification steps. The libraries were quantified and quality-checked using the Qubit (Thermo Fisher) and Bioanalyzer (Agilent). Paired-end 250 nt reads were generated using the MiSeq V2 500 cycle or V3 600 cycle kits (Illumina).

### Untargeted metagenomic Illumina sequencing

We followed the general outline of a previously developed protocol for untargeted sequencing of the mixed viral populations [34]. In brief, cDNA was generated as described for the amplicon-based methods. Second-strand cDNA was generated using *E. coli* DNA ligase and polymerase (New England Biolabs). The cDNA was purified by Agencourt AMPure XP beads (Beckman Coulter) prior to library preparation using Nextera XT (Illumina) following the vendor’s protocols, but with less reagents. Specifically, for tagmentation (12.5 μL reaction), we concentrated our cDNA to 4 μL using a DNA speedvac and used 5 μL of Tagment DNA Buffer (½ recommended) and 1 μL of Amplicon Tagment Mix (⅕ recommended). After incubation, the reaction was stopped using 2.5 μL of Neutralize Tagment Buffer (½ recommended). The libraries were indexed and amplified using ½ of the Nextera PCR reagents and primers in a 25 μL reaction. Agencourt AMPure XP beads (Beckman Coulter) were used for the final purification step (purified twice at a ratio of 0.7:1 beads to sample). The libraries were quantified and quality-checked using the Qubit (Thermo Fisher) and Bioanalyzer (Agilent). Paired-end 251 nt reads were generated using the MiSeq V2 500 cycle kit.

### Illumina data processing and variant calling using iVar

We developed an open source software package to process virus sequencing data and call iSNVs from technical replicates, iVar (intrahost variant analysis from replicates), and detailed documentation can be found at github.com/andersen-lab/ivar. A list of iVar commands and their brief descriptions are provided in **Table S5**, and additional details about the options are available in the documentation and can also be accessed in the help menu distributed with the tool (github.com/andersen-lab/ivar). iVar was used to write two pipelines for calling iSNVs from samples with or without the known reference sequence (*i.e.*, experimental and field-collected samples, respectively; **Fig. 9**). The software package was written in C++ and has two dependencies - HTSlib (github.com/samtools/htslib) and Awk (cs.princeton.edu/~bwk/btl.mirror/). Awk is generally available on most unix based operating systems and HTSlib has only one dependency - zlib (zlib.net/). The output of `mpileup` command from the widely used SAMtools [70] was used by the package to call variants and generate a consensus sequence from an alignment. The computational pipeline was divided in four main sections: (***1***) alignment, (***2***) trimming, (***3***) constructing nucleotide matrices and scan for iSNVs within primer regions, and (***4***) statistical comparisons between replicate datasets. The paired-end reads were aligned to a provided reference genome using BWA [71]. The primer sequences were trimmed from the reads using a BED file, with the primer positions, followed by quality trimming. iSNVs above the frequency cut-off of 3% were then called using the `mpileup` command from SAMtools. In addition to the frequency threshold, Fisher’s exact test was used to determine if the frequency of the variant is significantly higher than the mean error rate at that position (see contingency table, **Table S6**). Based on the iSNVs called, mismatches in the primer sequences were identified and the reads from the amplicon were removed. The variant calling step were repeated to remove any influence of the primer mismatch on iSNV frequencies.

**Figure 9.**
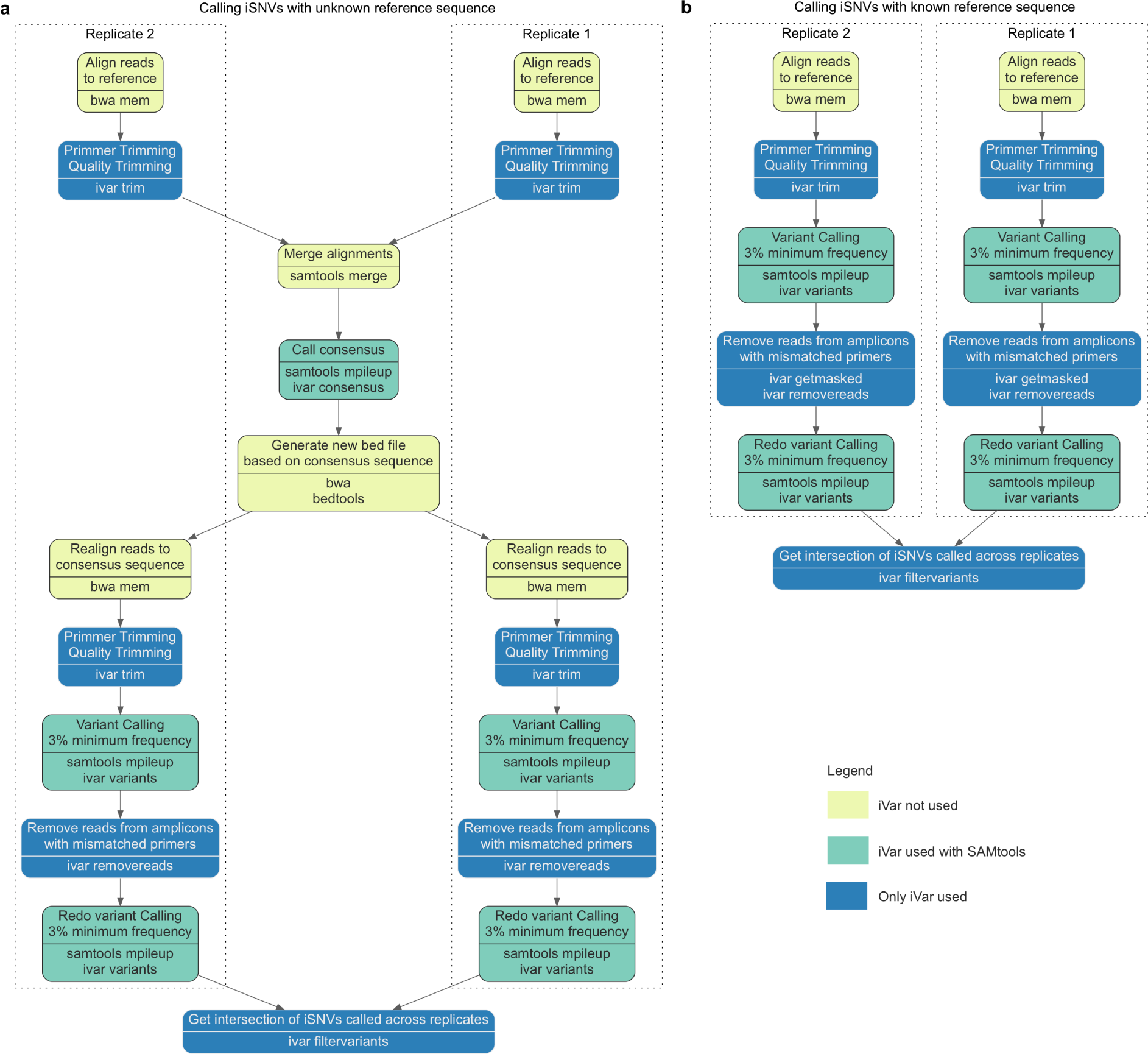
Overview of iVar pipeline. iVar contains two pipelines for calling intrahost single nucleotide variants (iSNVs) from samples (**a**) without and (**b**) with the known reference sequence. The nodes in the chart are colored based on the usage of iVar, SAMtools, bwa, and SAMtools at each step. (**a**) For samples with a known reference sequence, the primer sequences are trimmed from the sequenced reads, followed by quality trimming. The iSNVs are then called with a minimum frequency threshold of 3%. Primers with mismatches are then identified and the reads corresponding to the mismatched primers are removed from the alignment to ensure that any bias introduced in the iSNV frequencies is removed. The variant calling step is repeated on the filtered alignment file. The steps are repeated across all the replicates of the sample and an intersection of the iSNVs across all the replicates are considered to be the “true” iSNVs. (**b**) For samples with an unknown reference sequence, the iSNVs cannot be called directly using a reference sequence. In this case, an alignment to a reference sequence in done after quality and primer trimming for all the replicates. The resulting alignment BAM files are merged and a majority consensus sequence is called. Reads are aligned back to this consensus sequence, for each replicate and iSNVs are called using this new alignment at a minimum frequency of 3%. The reads corresponding to mismatched primers are then removed and iSNVs are called across all replicates. An intersection of iSNVs across all replicates are considered to be the “true” iSNVs.

### Validation of iVar

iVar was validated against existing tools. (**Table S7, Fig. S1-S3**). We validated iSNV calling in iVar against the `mpileup2snp` and `mpileup2indel` commands in VarScan2 (v2.3.9) [43] using four datasets - two simulated datasets and two clinical Zika virus samples, sequenced using PrimalSeq. We ran both tools with no thresholds and with a quality threshold of 20 and a minimum frequency threshold of 3%. We validated the consensus calling in iVar against the consensus calling available in Geneious (v11.1.4) [45] using four datasets - two simulated datasets and two clinical Zika virus samples, sequenced using PrimalSeq. We did the consensus calling at four different thresholds - 0%(majority), 25%, 50%(strict) and 90%. We counted the mismatches between the resulting consensus sequences from iVar, Geneious and the reference sequence by performing a multiple sequence alignment using MAFFT (v7.388) [44]. While counting mismatches, we ignored mismatches when one sequence had a gap and the other had a ‘N’, since in either case. We validated primer trimming and quality trimming in iVar against anchored adapter trimming in cutadapt (v1.16) [47] and against Trimmomatic [46]. iVar uses a sliding window starting from the 5’ end and checks if the average quality within the window drops below the threshold. As soon as the quality drops below the threshold, it trims the sequence by soft clipping the read. This is different from the algorithm cutadapt uses to do quality trimming, but is similar to the sliding window approach used by Trimmomatic. The data and code used for the validation are at github.com/andersen-lab/ivar-validation/.

### Oxford Nanopore sequencing and analysis

Using the same PCR amplicons used for amplicon-based Illumina sequencing, we sequenced three replicates of the mixed Zika virus population (90% virus #1, 10% virus #2) using the Oxford Nanopore GridION sequencer. Native, 1D barcode libraries (SQK-NSK007, Oxford Nanopore Technologies, UK) were prepared according to previously published methods [17], with three amplification replicates corresponding to barcodes 1, 2 and 3. The pooled sequencing library was sequenced on an R9.4 version flowcell (FLO-MIN106, Oxford Nanopore Technologies, UK). Reads were basecalled using Albacore 2.3.1 using the command-line read_fast5_basecaller.py -c r94_450bps_linear.cfg -i fast5 -o fastq -r -t 12. Reads were subsequently demultiplexed with Porechop 0.2.3_seqan2.1.1 using default (lenient) settings (github.com/rrwick/Porechop). A total of 2.4 million reads were generated which after alignment and trimming covered 95.66% of the reference genome (Genbank KX087101). For the purposes of assigning to genotypes (*i.e.*, unique virus haplotypes), reads were assigned to individual strains using BWA-MEM [72] against a custom reference database comprising four Zika virus genomes: Genbank KX087101 (virus #1), KU955593 (virus #2), EU545988 (an Asian lineage virus isolated in 2007) and NC_012532 (MR766, an African lineage virus). Counts for each assignment were retrieved, ignoring multi-mapping reads using the shell command bwa mem -x ont2d | samtools view -h -F 256 - | samtools view -h -F 2048 - | cut -f 3 | sort | uniq -c. Next, each replicate was aligned to the PRVACB59 reference genome with BWA-MEM using setting -x ont2d. Primer binding sites and any residual adaptor sequence were masked in the resulting BAM alignment using the align_trim script from the Zibra pipeline [17]. Allele frequencies and putative iSNVs (ignoring insertions or deletions) were extracted from BAM files using a Python script freqs.py (included in the accompanying code repository: github.com/nickloman/zika-isnv). This script utilises the pileup functionality of samtools via the pysam Python interface module (github.com/pysam-developers/pysam). Only predicted variants with more than 10 supporting forward and 10 supporting reverse reads were considered. The logistic regression model was trained and tested under a ten-fold cross validation scheme using the train function with the parameters method=“glm” and family="binomial” from the caret (github.com/topepo/caret/) library in R. Class probabilities for the ROC curve were captured from the same function and plotted using ggplot2 (github.com/tidyverse/ggplot2).

### Diversity metrics

Virus iSNVs >3% were used to genetically characterize the populations derived from *in vitro*, *in vivo*, and field samples (**Fig. 6 & 7**). For this purpose, insertions and deletions were not analyzed. For the Zika virus *in vitro* and *in vivo* samples, where the ancestral PRVABC59 sequence was known, the iSNV frequencies were calculated by the difference between the ancestral (pre-infection) and derived (post-infection) frequencies (*e.g*., if variant X was detected at 5% in the ancestral and 10% in the derived population, the iSNV frequency was listed as 5%). For the Zika and West Nile virus field samples, where the ancestral virus sequence was not known, the derived iSNV frequencies were used for the population genetic analysis. Richness was calculated by the total number of iSNV sites per population (**Fig. 7a**). Complexity, uncertainty associated with randomly sampling an allele, was calculated at each site using Shannon entropy: 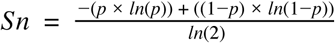, where *p* is the iSNV frequency and the mean *Sn* from all evaluated sites within the virus genome was used to determine the population complexity (**Fig. 7b**). Distance was calculated by the sum of all of the iSNV frequencies per population (**Fig. 7c**). Selection was estimated by the proportion of nonsynonymous variants: 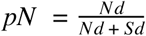, where *Nd* is the sum of the nonsynonymous iSNVs and *Sd* is the sum of the synonymous iSNVs within the population (**Fig. 7d**). Synonymous and nonsynonymous iSNVs were identified using custom python scripts that compare codons from the consensus and minor alleles.

## Additional files

All additional files can be found at github.com/andersen-lab/paper_2018_primalseq-ivar. A cookbook is also included to run iVar for two replicate Zika sequencing runs.

**Additional file 1: Table S1.** Location of PCR primer mismatches to divergent Zika viruses. **Table S2.** Laboratory and field-collected Zika and West Nile virus samples used in this study and sequencing statistics. **Table S3.** Primer sequences for tiled amplification of Zika virus. **Table S4.** Primer sequences for tiled amplification of West Nile virus. **Table S5**. A list of commands and descriptions available in iVar. **Table S6.** Contingency table for Fisher’s exact test to determine if the frequency of the variant is significantly higher than the mean error. **Table S7**. The list of commands in iVar and the tool that was used for validation, (Excel).

**Additional file 2:** Laboratory protocol for generating sequencing libraries for measuring intrahost virus genetic diversity. (PDF).

**Additional file 3: Supplemental Figures S1-S3**. Validation of iVar for intrahost single nucleotide variant calling (**Figure S1**), consensus calling (**Figure S2**), and trimming (**Figure S3**). (PDF).

## Declarations

### Availability of data and materials

All additional files can be found at github.com/andersen-lab/paper_2018_primalseq-ivar. The laboratory and computational (iVar) protocols generated from this study can be found in **Additional file 1**, **Additional file 2**, and github.com/andersen-lab/ivar. Protocol updates and additional primer schemes can be found at grubaughlab.com/open-science/amplicon-sequencing/ and andersen-lab.com/secrets/protocols/. The validation analyses from this study can be found in **Additional file 3**, github.com/andersen-lab/ivar-validation/, github.com/nickloman/zika-isnv, and NCBI Bioproject PRJNA438514.

### Authors’ contributions

The study was conceived and coordinated by NDG, KG, NJL, and KGA. The samples were provided by BJM, ALT, LMP, DEB, SG, NG, KKAVR, SI, SFM, and LLC. Library preparation and sequencing was performed by NDG, JGDJ, and JQ. The variant calling pipeline (iVar) was designed and built by KG and NM. The data was analyzed and interpreted by NDG, KG, NJL, and KGA. The manuscript was written by NDG, KG, NJL, and KGA with input from all co-authors. All authors read and approved the final manuscript.

#### Acknowledgements

We thank Barney Graham (VRC; NIAID/NIH) for support with the non-human primate studies, the Florida Department of Health for providing clinical samples, Miami-Dade Mosquito Control for providing collected *Ae. aegypti* pools, San Diego County Vector Control Program for providing West Nile virus samples, and Glenn Oliveira, Mark Zeller, Refugio Robles, Emily Spencer, Dylan Grubaugh, and Sophie Taylor for technical support.

